# Trends in genome diversity of small populations under a conservation program: a case study of two French chicken breeds

**DOI:** 10.1101/2024.02.22.581528

**Authors:** Chiara Bortoluzzi, Gwendal Restoux, Romuald Rouger, Benoit Desnoues, Florence Petitjean, Mirte Bosse, Michèle Tixier-Boichard

## Abstract

Livestock biodiversity is declining globally at rates unprecedented in human history. Of all avian species, chickens are among the most affected ones because many local breeds have a small effective population size that makes them more susceptible to demographic and genetic stochasticity. The maintenance of genetic diversity and control over genetic drift and inbreeding by conservation programs are fundamental to ensure the long-term survival and adaptive potential of a breed. However, while the benefits of a conservation program are well understood, they are often overlooked. We here used temporal whole-genome sequencing data to assess the effects of a conservation program on the genetic diversity (Δ*π*), deleterious variation (ΔL), and inbreeding (ΔF) of two local French chicken breeds, the Barbezieux and Gasconne. We showed that when the conservation program is consistent over time and does not undergo any major organisational changes (i.e., Barbezieux), the loss of genetic diversity is limited. This was true for both pedigree and genomic inbreeding but also for the genetic load estimated from functionally important genome-wide variants. However, when a conservation program is interrupted or re-initiated from scratch (i.e., Gasconne), the loss of genetic diversity can hardly be limited as a result of the bottleneck effect associated with the re-sampling. Our results reinforce the imperative to establish and sustain existing conservation programs that aim to keep populations with a relatively small effective population size from the brink of extinction. Moreover, we conclude by encouraging the use of molecular data to more effectively monitor inbreeding at the genome level while improving fitness by tracking protein-coding and non-coding deleterious variants.

## Introduction

Livestock breeds are recognised as important components of world biodiversity since they harbor genetic variants that can be useful to agriculture in the future. Nevertheless, livestock diversity is declining globally, as shown by the high rate of world’s livestock breeds reported in The State of the World’s Biodiversity for Food and Agriculture of the Food and Agriculture Organization (FAO) as being at risk of extinction (Scherf, Pilling, et al., 2015). Among the 7,745 worldwide local livestock populations, 26% are at risk of extinction, 67% are of unknown risk status, and 7% are classified as extinct (Bélanger, Pilling, et al., 2019). Europe and the Caucasus region have the highest number of extinct mammalian and avian breeds than any other region, reaching values of 78% and 86%, respectively.

Avian species, and particularly chicken, are among the livestock species with the highest percentage of breeds with a critical status, although difference can be observed at the national and regional level; for instance, in France, of the 47 local poultry breeds, 46 have the status of endangered, as highlighted in a recent report of the French Ministry of Agriculture (Bouffartigue et al., 2023). The establishment in the mid 20th century of few, specialised breeding industries that rely on a few selected lines for egg (layer) or meat (broiler) production has been partially responsible for the decline in local chicken diversity in Europe and North America (Muir et al., 2008). However, the large number of chicken breeds at risk is also due to the often unclear and problematic definition of a breed, which makes any direct risk assessment rather challenging. From a genetic perspective, local chicken breeds are at major risk of extinction because their small population size makes them more susceptible to stochastic demographic and genetic events. The risk of genetic erosion is often enhanced by the lack of conservation programs, either on farm, by livestock breeders in the production system (i.e., *in situ*) or in dedicated facilities, such as ark farms or experimental facilities (i.e., *ex situ in vivo*) (Bortoluzzi, Crooijmans, et al., 2018). Furthermore, semen cryopreservation (i.e., *ex situ in vitro* conservation) is still not routinely implemented in chickens.

Genetic drift, or the random fluctuation in allele frequencies, is the main stochastic event responsible for the loss of genetic diversity in small populations (Fernández, Meuwissen, et al., 2011). In fact, genetic drift can reduce the viability and adaptive potential of a population. Recent studies in wild and domesticated species (Abascal et al., 2016; Bortoluzzi, Bosse, et al., 2020; Robinson et al., 2019; Van Der Valk et al., 2019; Xue et al., 2015) have shown that the risk of extinction in small populations is also a consequence of harmful mutations that lower the fitness of an individual carrying them. The rationale is the reduced efficiency of natural selection at purging harmful mutations because of genetic drift (Kimura, 1957; Ohta, 1973). Therefore, harmful mutations can accumulate and reach fixation in the genome. Additionally, as small populations suffer from inbreeding resulting from mating between close relatives (Kardos et al., 2016), (recessive) mutations in homozygous state can express their harmful nature.

Conservation programs are able to maintain genetic diversity while controlling for genetic drift (De Cara et al., 2013; Fernández, Meuwissen, et al., 2011; Fernández, M Toro, et al., 2004). However, the impact of a conservation program on a population in terms of genetic diversity, deleterious variation, and inbreeding have rarely been investigated in local livestock breeds at the whole-genome level. Such assessment is of particular relevance today, as the maintenance of high genetic diversity alone is not sufficient to ensure the long-term survival of populations of small size (Oosterhout et al., 2022). Recent advances in sequencing technologies can help us in the task of evaluating a conservation program with the aim of providing objective recommendations to effective management practices for small local populations (Díez-del-Molino et al., 2018; Habel et al., 2014). Temporally sampled genomic data are a powerful tool to monitor changes in genetic parameters, including genetic diversity (Δ*π*), inbreeding level (ΔF), deleterious variation (ΔL), and, if applicable, selection (ΔS), as illustrated in the case of the Spanish cattle breed Asturiana de Los Valles (Boitard et al., 2021). Hence, when possible, temporal genomic indices should be quantified to evaluate and guide existing and future conservation programs (Díez-del-Molino et al., 2018).

In this study, we assessed the impact of 10 generations of a conservation program on the genetic and deleterious variation of two local French chicken breeds, the Barbezieux and Gasconne, by means of whole-genome sequencing data. For each breed, a conservation program was established in 2003 by the breeders’ association in collaboration with a professional breeding center *The Centre de Sélection de Béchanne*, with the methodological support of the French Union of Poultry and Fish Breeders (SYSAAF) for the management of pedigree data and mating plans. However, while the conservation program of the Barbezieux continued with a one-year generation interval, that of the Gasconne was discontinued and completely replaced in 2009 with a new set of founder sires and dams, unrelated to those used in 2003. Yet, the common point between these two breeds was the very small number of founders, being less than 10 sires and 10 dams.

To assess the effectiveness of these two conservation programs at maintaining genetic variation, temporal genomic erosion between 2003 and 2013 was analysed by quantifying delta indices related to genetic diversity (Δ*π*), inbreeding (ΔF), and deleterious variation (ΔL), which were ultimately used as reference to provide recommendations for future management practices.

## Material and methods

### Sampling statement

Data used in this study were collected as part of routine data recording for a conservation program. Blood samples collected for DNA extraction were conducted under veterinary care for routine health monitoring and only used for the conservation program, in line with the French law on the protection of farm animals.

### History of the populations

Two local chicken breeds, the Barbezieux and Gasconne, were chosen for this study because of their management history and availability of gene bank samples at two time periods. The origin of the two breeds dates back to the 19th century in South-west France in the city of Barbezieux-Saint-Hilaire for the Barbezieux and the city of Masseube for the Gasconne (Fig. 1a). Both breeds are considered as dual-purpose breeds, laying about 200 eggs per year while producing high quality meat. They are robust and generally raised in a free range system. They are generally valued locally in the short chain market where they benefit from a designation of origin.

**Figure 1.**
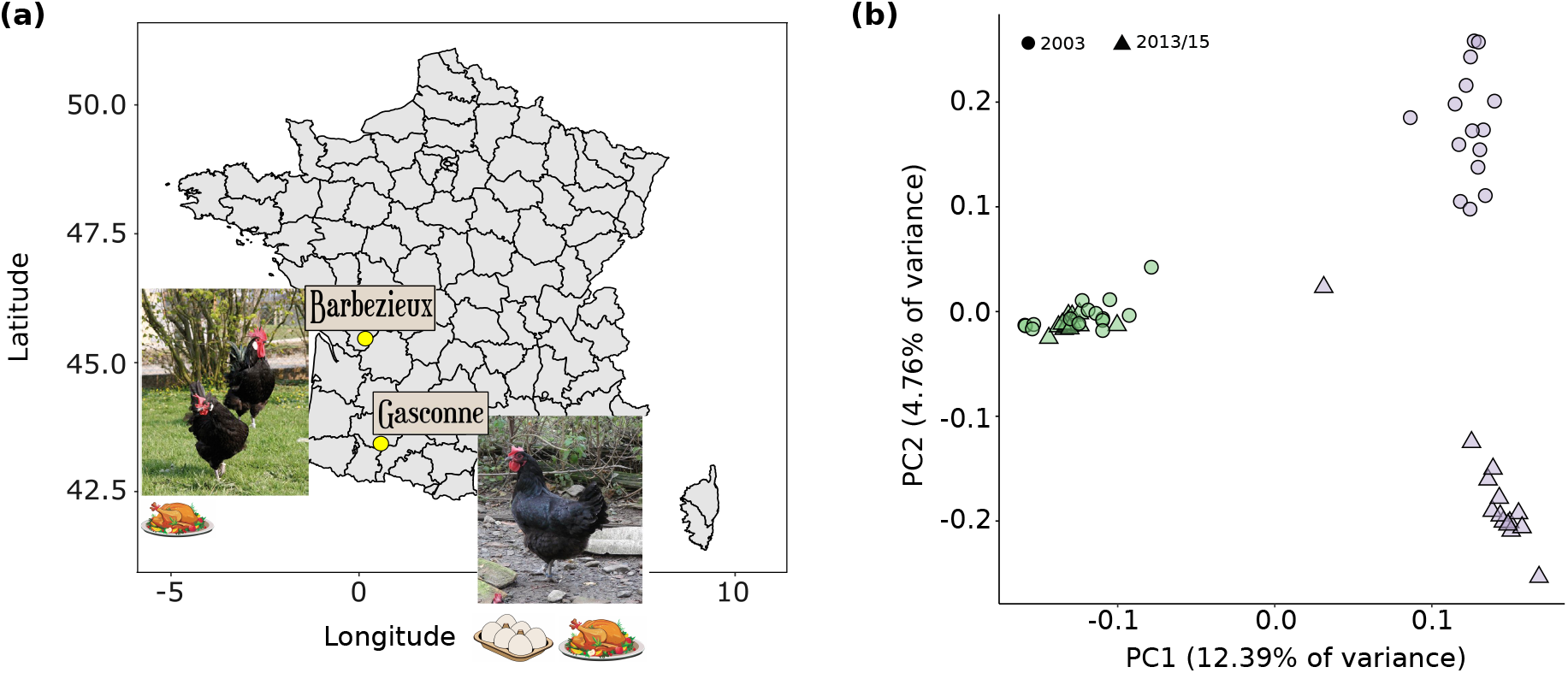
Samples and population structure. **a**. Geographic origin of the Barbezieux and Gasconne breed, with relative breeding objectives (meat or meat/egg). **b**. Principal component analysis (PCA) performed using 12,168,183 bi-allelic SNPs after filtering for a missing rate of 10% and a minor allele frequency (MAF) of 0.05. Individuals from the Barbezieux breed are coloured in green, while those from the Gasconne breed are in purple. Individuals belonging to the generation sampled in 2003 are represented by circles, while those from the generation sampled in 2013/15 are represented by triangles. The Gasconne individual sampled in 2013 that is found far from the remaining samples is individual 7218.

In 2003, both breeds were included in a research project aimed at defining the main parameters which determine the success and sustainability of exploitation programs for local breeds (Tixier-Boichard et al., 2006). The project started simultaneously for both breeds with the first animals born in 2003 recorded in a pedigree at the Breeding Center of Bechanne for the Barbezieux, with 14 founder sires and dams, and at the Agricultural school of Saint Christophe for the Gasconne, with 25 founder sires. For each breed, a breeders’ association was set up to define the breeding objectives and to monitor the management program. Thanks to the DNA bank established in the frame of the French Center for Animal Biological Resources, CRB-Anim research infrastructure (https://doi.org/10.15454/1.5613785622827378e12), we had access to samples for an equal number of individuals, for each breed, both at the start of the conservation program (i.e., founder population) and 10 generations after to get a good picture of each population (Table 1).

**Table 1.**
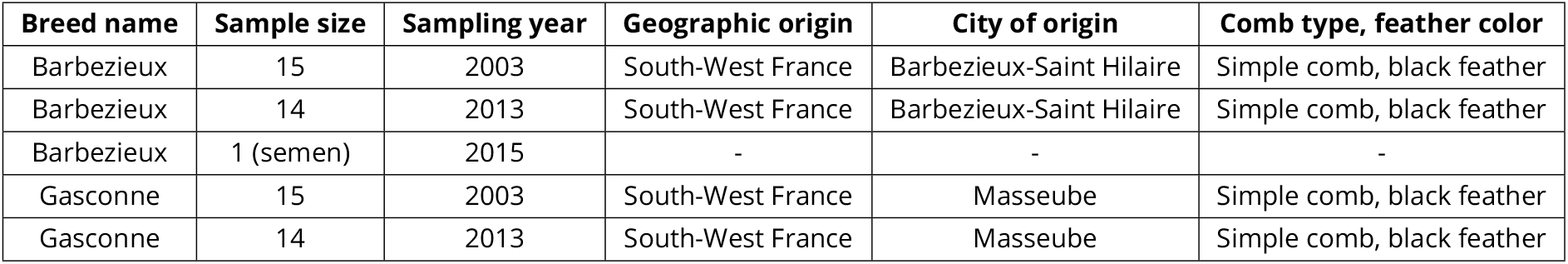
Samples sequenced for this study.

### Sampling

To perform a time series analysis and to monitor the impact of management practices across 10 generations, we sampled 15 founder individuals born in 2003 for each breed. Then we completed these samples with 14 individuals born in 2013 for both Barbezieux and Gasconne breeds (Table 1). In addition, we completed the Barbezieux breed sampling with the semen of one male collected in 2015, bringing the total sample size to 59. Except for this latter one, all samples consisted of DNA extracted from blood. Siblings and half siblings were discarded from the selection process to minimise relatedness in the dataset. For each breed we also obtained the following additional information: (1) complete pedigree data for the period 2002-2019 for the Barbezieux and 2009-2019 for the Gasconne; (2) body weight at 8 weeks of age from 2003 to 2019 for the Barbezieux and from 2010 to 2019 for the Gasconne; and (3) six reproductive traits for the period 2003-2018 and 2011-2018 for the Barbezieux and Gasconne, respectively, defined as the average number of eggs set in the incubation, the average number of infertile eggs, average number of hatched eggs, % fertile eggs, % hatched eggs, and late embryonic mortality.

Pedigree and phenotypic data were provided by the SYSAAF under a data transfer agreement signed with the breeders’ association for each breed.

### Sequencing, read processing and alignment

Sequencing was carried out on a NovaSeq 6000 sequencing machine using standard double-stranded library preparation protocols. Sequencing statistics are given for each individual in Table S1 of the Supplementary Material. All analyses were based on an alignment of sequence data from all samples to the chicken GRCg7b reference genome (GenBank assembly accession: GCA_016699485.1). Sequence data were mapped to the chicken reference genome with the BWA-mem2 v2.2.1 algorithm (Vasimuddin et al., 2019) using default options. Aligned reads were sorted using samtools v1.9 (Danecek et al., 2011), while duplicate reads were tagged and removed using Picard v2.26.2 (https://broadinstitute.github.io/picard/). Base quality recalibration was carried out in GATK v4.2.4.0 (McKenna et al., 2010) using known variants downloaded from the Ensembl Release 112 (https://ftp.ensembl.org/pub/release-112/variation/vcf/gallus_gallus/), which were imported from dbSNP and remapped to the chicken GRCg7b. SNPs and InDels were called simultaneously in each sample via local *de-novo* assembly of haplotypes using the *HaplotyeCaller* tool in GATK. Individual GVCF files were merged into a single GVCF file using the *CombineGVCFs* tool to perform joint genotyping on all samples using the *GenotypeGVCFs* tool, while retaining variants with a mapping quality >30 and base quality >10. Commonly called variants were further filtered using the *VariantFiltration* tool following GATK recommendations in order to have a more accurate call set. We performed an additional filtering step on the final VCF file using a custom python script (see **Data availability**), retaining genotypes whose coverage was between 4x and 2.5 the individual mean genome-wide coverage estimated with samtools depth.

### Principal component analysis

A principal component analysis (PCA) of genetic variation was carried out in SNPRelate (Zheng et al., 2012) for R v4.3.2 to detect any existing structure within and between the two breeds. For this analysis, we used the *snpgdsPCA* command on autosomal SNPs only (autosome.only=TRUE), after removing monomorphic SNPs (remove.monosnp=TRUE). The first PCA was performed on all samples, considering as input only bi-allelic SNPs with a missing rate *<*10% (missing.rate=10) and a minor allele frequency of 0.05 (*n* = 12,168,183 SNPs). We did not perform any linkage disequilibrium (LD) pruning in SNPRelate to avoid excluding sites corresponding to fixed differences between the two breeds. In addition to the all-samples PCA, we performed a breed-specific PCA, in which bi-allelic SNPs were also pruned in SNPRelate using the *snpgdsLDpruning* command for an |LD| threshold of 0.5. After pruning, 65,665 and 88,769 SNPs remained for the Barbezieux and Gasconne, respectively. Population differentiation was further analysed by estimating the fixation index (*F*_*st*_) between populations (i.e., combination of breeds and time period) in consecutive non-overlapping 50-kb windows in VCFTools v0.1.16 (Danecek et al., 2011) after removing windows with less than 300 SNPs.

We also built a Neighbor-joining (NJ) phylogenetic tree in Phylip v3.697 (https://phylipweb.github.io/phylip/) from the identity-by-state (IBS) distance relationship matrix estimated in Plink v1.90b6.2.1 (Purcell et al., 2007) on 12,476,884 SNPs after filtering for a missing rate *<*10% and a minor allele frequency (MAF) *<*0.05..

### Genome-wide heterozygosity

Heterozygosity was calculated for each individual separately as the number of heterozygous genotypes in consecutive non-overlapping windows of 100-kb using a custom python script (see **Data availability**). Heterozygosity was calculated for the entire autosomal genome (InDels excluded) and is here expressed as the number of heterozygous sites corrected for the number of sites that did not meet the coverage criteria (i.e., 4x ≤ coverage ≤ 2.5*mean genome-wide coverage) and the window size (i.e., 100,000 bps) (Bortoluzzi, Bosse, et al., 2020; Bosse, Megens, Madsen, Paudel, et al., 2012). However, only windows where at least 80% of the sites met the coverage criteria were considered for the individual genome-wide heterozygosity (Bortoluzzi, Bosse, et al., 2020; Bosse, Megens, Madsen, Paudel, et al., 2012).

### Within-individual runs of homozygosity

Runs of homozygosity (ROH), here defined as genomic regions showing lower heterozygosity than expected based on the average genome-wide heterozygosity, were identified using the approach of Bortoluzzi, Bosse, et al. (2020) based on Bosse, Megens, Madsen, Paudel, et al. (2012). To identify ROH, we first calculated the corrected number of heterozygous genotypes in consecutive non-overlapping 10-kb windows along the genome of each individual. For this step, we considered only 10-kb windows where at least 80% of the sites met our coverage criteria. We then considered ten consecutive 10-kb windows at a time (i.e., 100-kb) and applied two filtering steps. First, we calculated the level of heterozygosity within the 10 consecutive windows - here indicated as *π*w - and retained only those for which *π*w was below 0.25 the average genome-wide heterozygosity - here indicated as *π*g. We used a threshold of 0.25 as this value was found to be able to filter out windows enriched for heterozygous sites. In the second step we tried to reduce the impact of local assembly and alignment errors as much as possible by relaxing another set of parameters within the retained 10 consecutive windows - from here onwards we will refer to these windows as candidate homozygous stretches.

Sequence data are prone to assembly and alignment errors and very often these errors result in a peak of heterozygous sites. To filter out these peaks, we first looked at each window making up the candidate homozygous stretch to identify any window whose heterozygosity was twice that of the genome (*π*g). If *π*w did not exceed 20% the average genome-wide heterozygosity also when considering windows with a peak in heterozygosity, then the candidate homozygous stretch was retained. Otherwise, the candidate homozygous stretch was splitted into smaller stretches. ROH were finally classified - based on their size - into short (100 Kb - 1 Mb), medium (1 - 3 Mb), and long (≥ 3 Mb).

### Between-individual sequence identity

To identify genomic regions shared between individuals (identity-by-descent segments or IBD), we first resolved the phase of the distinct haplotypes within each sample using Beagle v5.3 (B Browning, Tian, et al., 2021). Following (Wu et al., 2023), phasing was performed on the all-samples dataset of filtered variants using 10 burnin iterations, 12 phasing iterations, a window length of 0.02 cM, a window overlap of 0.01 cM, and an effective population size of 100,000 (Wu et al., 2023). Phasing was performed on each chromosome separately. IBD segments between individuals were identified using the following parameters in the refinedIBD program (B Browning and SR Browning, 2013): 0.06 cM window length, a minimum length of 0.02 cM to report an IBD segment, a trim value of 0.001 cM, and a LOD score of 3.0 (Wu et al., 2023).

### Pedigree- and genomic-based inbreeding

We used the pedigree provided by the SYSAAF to estimate the pedigree inbreeding coefficient, *F*_*ped*_, in 43 of the 59 samples. Samples that were not present in the pedigree and were thus excluded from the *F*_*ped*_ estimation were the 15 Gasconne founders and the Barbezieux sample from 2015. The pedigree inbreeding coefficient was calculated using the *calcInbreeding* function of the pedigree library v1.4.2 (https://rdocumentation.org/packages/pedigree/versions/1.4.2) for R v4.3.2. In addition to the expected inbreeding, we used our set of ROH to estimate the realised inbreeding, or *F*_*ROH*_, here expressed as the ratio between the total length of ROH within an individual (*L*_*ROH*_) and the length of the autosomal genome (*L*_*auto*_ = 945,968,431 nucleotides) (McQuillan et al., 2008). We did not exclude complex regions, as duplicate reads were removed as much as possible in the alignment step (see **Sequencing, read processing and alignment**). However, we did exclude from *L*_*auto*_ both sex chromosomes and the mitochondrial genome, as these were also excluded from the heterozygosity and ROH analyses.

### Liftover chain file generation

We generated a Liftover chain file between the chicken GRCg6a (GenBank accession: GCA_000002315.5) and chicken GRCg7b reference genome (GenBank accession: GCA_016699485.1). A repeat-masked (or soft-masked) version of both reference genomes was downloaded from Ensembl release 106 and 112, respectively. The Liftover chain file was generated following the UCSC approach. We first generated in Lastz v1.04.00 (Harris, 2007) a pairwise alignment between the old (GRCg6a) and new (GRCg7b) chicken reference genome, using the following parameters: –hspthresh=2200, –inner=2000, –ydrop=3400, –gappedthresh=10000, and the HoxD55 substitutions score matrix (G Zhang et al., 2014). The obtained alignments in axt format were chained together using *axtChain*, by specifying the –linearGap option to lose. We then ran *chainMergeSort* to combine all sorted chain files into a larger, sorted chain file, on which we ran *chainNet* and *netChainSubset* to remove chains that do not have a chance of being netted. The resulting Liftover chain file was used to liftover the genomic positions of the ch(icken) CADD scores (see **Annotation of variants**).

### Polarisation of variants

The bias towards the alleles present in the reference sequence (reference bias) can lead to inaccurate genomic analysis. To reduce the effect of the reference bias, we polarised all alleles present in our dataset as ancestral or derived using the 10 fowl reference-free multiple sequence alignment generated with cactus (Paten et al., 2011) and downloaded from Ensembl release 112 (https://ftp.ensembl.org/pub/misc/compara/multi/hal_files/Fowl-10-way_20240131.hal). For our polarisation approach, we used the reconstructed ancestral sequence of the 7 members of the Galliformes order (Figure S1). We retained only bi-allelic SNPs for which either the reference or alternative allele matched the ancestral allele, while ancestral alleles that did not match either chicken allele were discarded.

### Annotation of variants

Polarised variants were annotated using the Ensembl Variant Effect Predictor (VEP) (release 110) (McLaren et al., 2016) and the Combined Annotation-Dependent Depletion tool developed for chicken (chCADD) (Groß et al., 2020). The VEP was limited to the annotation of protein-coding variants, whereas chCADD was used to equally annotate all variants in an individual’s genome, independently of their coding potential. VEP was run offline by specifying “gallus_gallus” as species after caches were downloaded for the chicken GRCg7b reference genome from Ensembl release 110 (https://ftp.ensembl.org/pub/release-110/variation/vep/). Amino acid substitution effects on protein function were predicted with SIFT (Kumar et al., 2009), which is based on sequence homology and the physical properties of amino acids. Before classifying our variants into functional classes, we applied a combination of filtering steps to improve the reliability of the prediction using a combination of VCFTools v0.1.16 (Danecek et al., 2011) and custom python scripts (see **Data availability**). We retained (1) bi-allelic variants with a call rate of at least 70%; (2) variants outside repetitive elements as these genomic regions are often difficult to sequence and are thus prone to errors, (3) variants found in genes 1:1 ortholog between chicken and zebra finch to reduce the effect of off-site mapping of sequence reads, and (4) variants for which the RNA-seq expression coverage was of at least 200 in heart and liver tissues (https://ftp.ensembl.org/pub/rapid-release/species/Gallus_gallus/GCA_016699485.1/ensembl/rnaseq/) (Bortoluzzi, Bosse, et al., 2020; Derks et al., 2017).

### Functional classes

We considered protein-coding variants classified by the VEP as synonymous, missense tolerated (SIFT score >0.05), missense deleterious (SIFT score ≤0.05), and loss of function (LoF) (i.e., splice donor, splice acceptor, start lost, stop gained, and stop loss). To validate the set of variants belonging to the damaging group (deleterious missense mutations and LoF mutations), we assigned the Genomic Evolutionary Rate Profiling (GERP) score (Davydov et al., 2010) computed on the 27-sauropsids whole-genome alignment downloaded from Ensembl Release 112 (https://ftp.ensembl.org/pub/release-112/compara/conservation_scores/27_sauropsids.gerp conservation_score). The GERP score is a measure of sequence conservation across multiple species. Since conservation is often an indicator of strong purifying selection, GERP is an excellent predictor of fitness effects and variant deleteriousness (Huber et al., 2020). Hence, of the initial set of putative damaging mutations, only those with a GERP score >1.0 were considered truly damaging.

### Estimation of genetic load

Estimating an individual’s genetic load based on genomic data is challenging. We therefore expressed the genetic load using two different approaches. We initially expressed the genetic load as a function of the GERP score - here called GERP load - by considering, for each individual, only damaging mutations with a GERP score >1.0, after re-adapting the formula presented in Orlando and Librado (2019). Finally, we estimated the genetic load as a function of the chCADD score - chCADD load - by considering, for each individual, protein-coding and non-coding variants that belonged to functional classes with an average chCADD score >10. The chCADD load was calculated as:

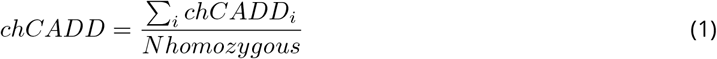

where *chCADD* is the score of a homozygous derived variant at genomic position i and *NHomozygous* is the total number of homozygous derived variants in each individual’s genome. Thus, the chCADD score measures variant deleteriousness and can effectively prioritise variants based on a comprehensive set of functional and evolutionary properties (Groß et al., 2020; Rentzsch et al., 2019).

### Signatures of selection

Genomic regions under positive selection were identified using the new generic Hidden Markov Model (HMM) developed by Paris et al. (2019). This HMM approximates the Wright-Fisher model implementing a Beta with spikes approximation, which combines discrete fixation probabilities with a continuous Beta distribution (Paris et al., 2019). The advantage of this model over existing ones is its applicability to time series genomic data. Prior to detecting regions under selection, we estimated the effective population size (*N*_*e*_) in each breed separately using the *currentNe* programme which can deal with specific domestic population features like small population sizes and family structures (Santiago et al., 2024). To estimate *N*_*e*_, we removed SNPs with an allele frequency *<*0.20 and >0.80 following recommendations (Paris et al., 2019) and selected a random number of 100,000 bi-allelic SNPs 100 times. The decision to randomly select SNPs was due to the fact that *currentNe* can process no more than 2 million SNPs. We then applied the HMM model developed by Paris et al. (2019) (https://pypi.org/project/selnetime), after which we removed SNPs with a false discovery rate (FDR) threshold of 5% as estimated in the q-value package (Storey et al., 2015) for R v3.2.0.

## Results

We generated whole-genome sequencing data from 30 Barbezieux and 29 Gasconne birds sampled between 2003 and 2015 (Table 1). All genomes were aligned, genotyped, and annotated with respect to the chicken GRCg7b reference genome, yielding a per-individual mean genome-wide depth >10x and mean mapping quality >30 (Table S1). Following variant calling and additional post-filtering steps, we identified 2 million InDels and 16 million SNPs distributed along the genome following a SNP density of 19.46/Kb (Table S2). We decided to limit our downstream analyses to the first 39 autosomes, excluding both sex chromosomes and the mitochondrial genome.

### Temporal changes in genetic diversity and inbreeding

The separation between the Barbezieux and Gasconne samples in the principal component analysis (PCA) confirms them as genetically distinct breeds (weighted *F*_*st*_: 0.108) (Fig. 1b; Figure S2). In the all-samples PCA and breed-specific PCA (Figure S3), we observed a clear differentiation between the Gasconne individuals sampled in 2003 and 2013 (weighted *F*_*st*_: 0.0595), which confirms a change of the population in the 10 generations period. The result was also confirmed by the Neighbor-Joining (NJ) analysis on the identity-by-state distance relationship matrix (Figure S4). By contrast, very little separation was observed between the two sets of birds sampled for the Barbezieux breed across the same period (weighted *F*_*st*_: 0.0160).

The analysis of genome-wide heterozygosity showed that, in 10 years, genetic diversity decreased by 2% and 10% in the Barbezieux and Gasconne, respectively, resulting in a genome-wide heterozygosity in 2013/2015 of *π*: 4.08×10-3 and *π*: 4.12×10-3, respectively (Fig. 2a). Temporal changes in genetic diversity (Δ) were significant in the Gasconne breed (Wilcox test *p-value*: 0.0157), but not in the Barbezieux breed (*p-value*: 0.2854).

**Figure 2.**
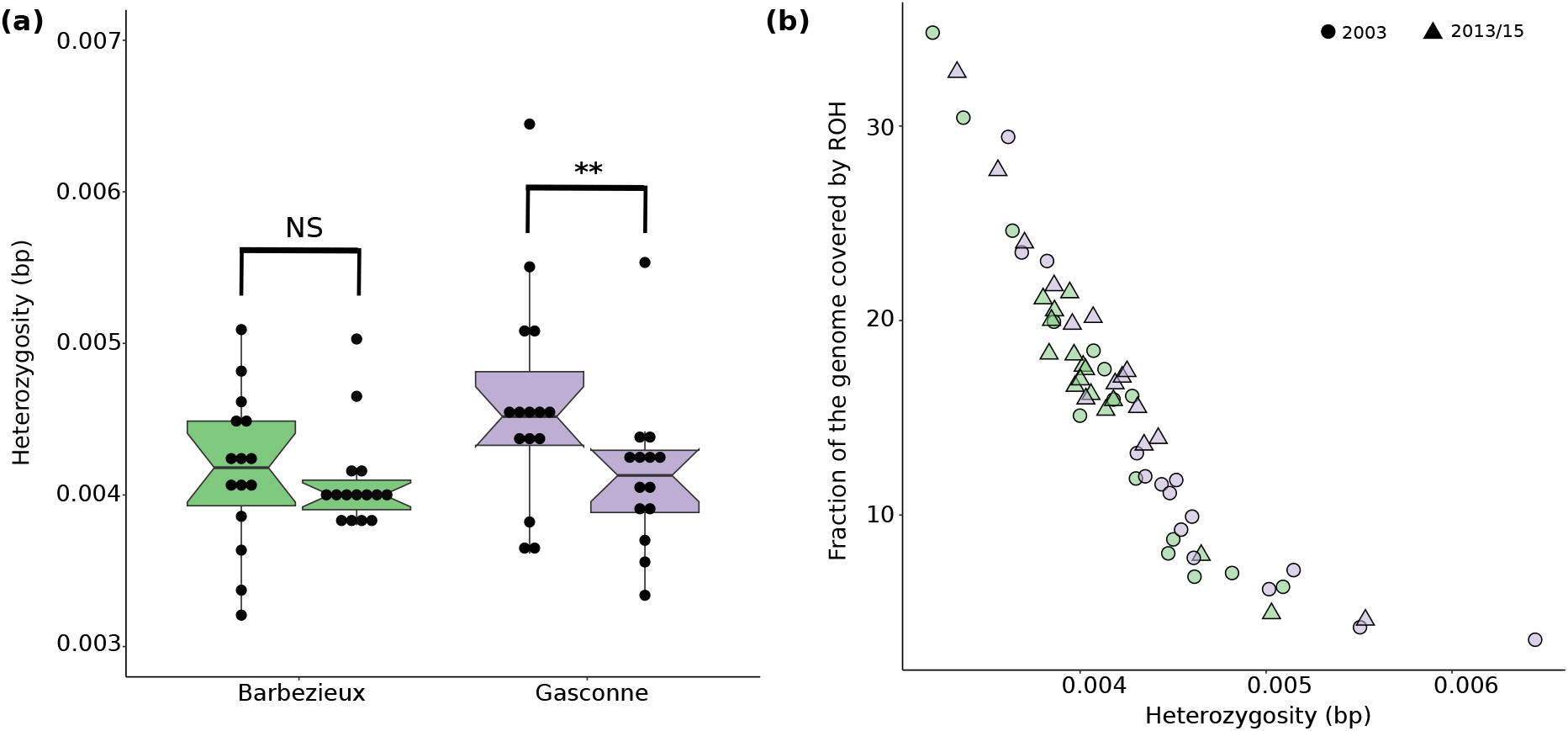
Temporal changes in genome-wide heterozygosity. **a**. Heterozygosity is the mean autosomal heterozygosity calculated for each individual and time point along the genome in consecutive 100 kb non-overlapping windows. **b**. Correlation between individual heterozygosity (bp) and fraction of the genome covered by runs of homozygosity (ROH).

Despite this faster decrease in heterozygosity, the Gasconne breed sampled in 2013/15 exhibited a higher within-breed diversity than the Barbezieux at the same sampling time. The within-breed reduction in genetic diversity observed in recent samples resulted from a fragmented heterozygosity distribution, where regions of high heterozygosity (Figure S5) were interspersed by regions enriched for homozygous genotypes, also defined as runs of homozygosity (ROH). Although mean genome-wide heterozygosity was negatively correlated with the total fraction of the genome covered by ROH (Pearson’s r: -0.90, *p-value*: *<*2.2×10-16) (Fig. 2b), the correlation did not capture the abundance and size distribution of ROH (Fig. 3a). Of all ROH size classes, long ROH (≥3 Mb) are of major concern as they result from recent close inbreeding. In Barbezieux individuals sampled in 2013/15 we identified a maximum of 15 long ROH, covering, on average, 4.42% of the genome, whereas in the Gasconne individuals we identified a maximum of 23 long ROH covering, on average, 5.96% of the genome (Table S3). In both breeds, the number of medium ROH increased over the 10 generations, whereas the number of long ROH increased only in the Gasconne breed (Figure S6), resulting in a larger fraction of the genome in homozygous state (18.66% versus 16.58% in the Barbezieux) (Fig. 3a; Figure S7). We also report a few ROH longer than 10 Mb, although these were mostly found in the Barbezieux individuals sampled in 2003. Of the individuals sampled in 2013/15, only 9 had no more than 2 ROH >10 Mb.

**Figure 3.**
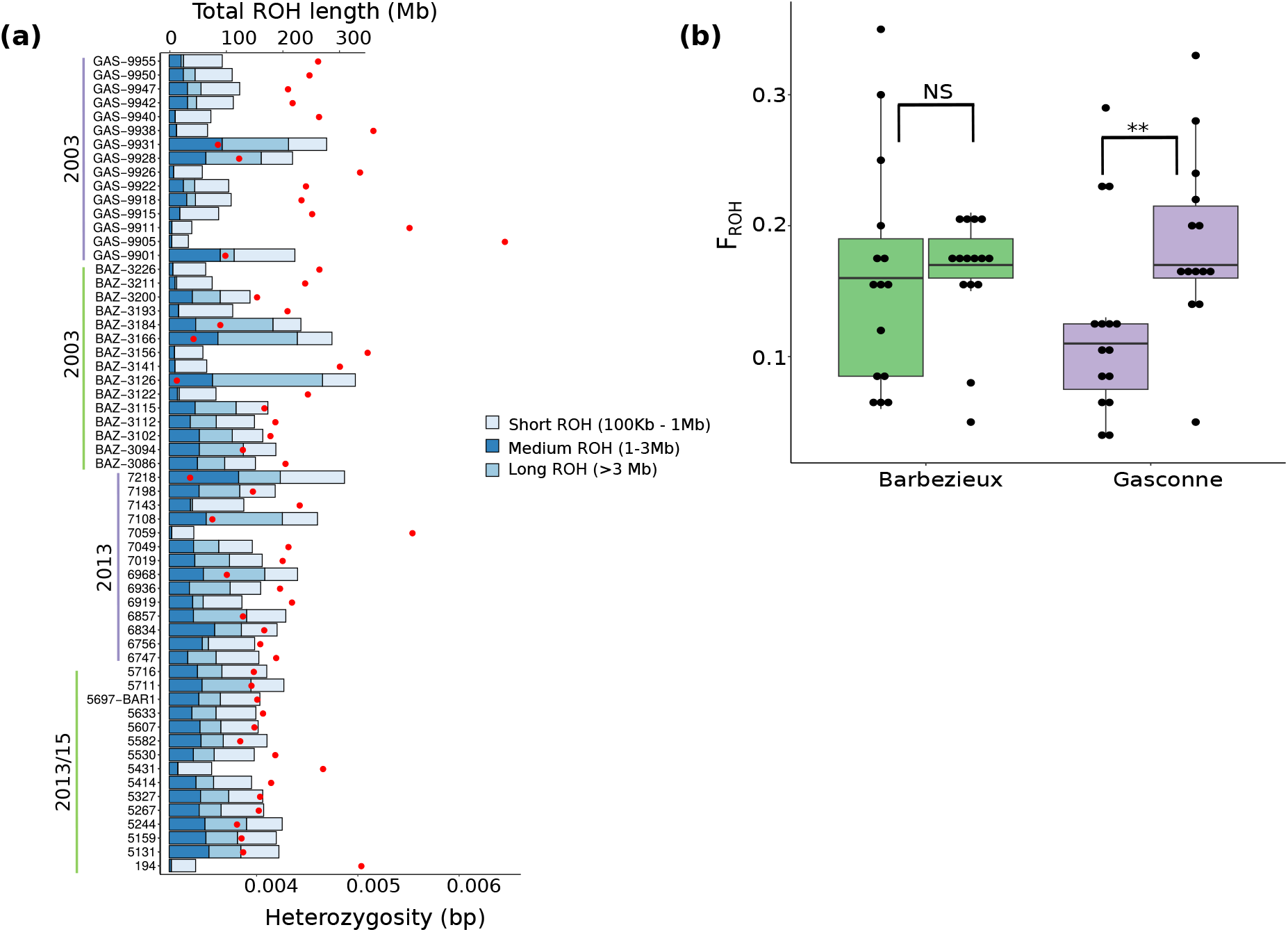
Runs of homozygosity (ROH) and temporal changes in inbreeding level. **a**. Average (autosomal) genome-wide heterozygosity per individual (red dot) and total length of short, medium, and long ROH per individual (barplot). **b**. Genomic (or realised) inbreeding coefficient estimated for each individual as a ratio between the total length of ROH within an individual and the actual length of the genome covered in our dataset.

For each breed we also examined haplotypes shared between individuals (identity-by-descent - IBD) as these provide information on the levels of recent inbreeding. To ensure a fair comparison with the ROH analysis, we retained only IBD segments >100Kb (or >0.1 cM). We found clear differences in the total length (in Mb) of IBD segments between breeds and time of sampling (Figure S8), consistent with the ROH analysis. In general, animals sampled in 2003 displayed longer IBD segments, although only a few shared segments longer than 90 Mb (Barbezieux) and 200 Mb (Gasconne), confirming the overall low relatedness among individuals sampled for this study. Overall, Gasconne individuals were found to share longer IBD segments than Barbezieux individuals, and this trend did not change over time, confirming the impact of recent inbreeding in this breed due to the change in management. Contrary, we observed a reduction in IBD size over time in the Barbezieux breed (Figure S8).

The genomic, or realised, inbreeding coefficient (*F*_*ROH*_) exhibited a 6.25% and 58.3% increase over time in the Barbezieux and Gasconne breed, respectively (Fig. 3b), although such increase was significant only in the Gasconne (Wilcox test *p-value*: 0.0092). We further calculated the pedigree inbreeding coefficient (*F*_*ped*_) (Table S4) and estimated the accuracy of *F*_*ped*_ in capturing individuals’ relationships. Although we were able to calculate the pedigree-based inbreeding for 29 Barbezieux and 14 Gasconne samples, we found that values of *F*_*ped*_ were much more homogeneous in the Barbezieux (*F*_*ped*_: 7%) than in the Gasconne (*F*_*ped*_: 3-15%), confirming the trend in *F*_*ROH*_ (Fig. 3b; Figure S9). We further correlated *F*_*ROH*_ and *F*_*ped*_ to verify the usefulness in a conservation program of the pedigree information. As expected, we report a significant positive correlation (Pearson’s r: 0.55; *p-value*: 1.42×10-4). The pedigree provided by the SYSAAF was also used to quantify changes in the number of sires and dams over the 10 generations (Figure S10). The conservation program was able to increase the number of breeding males and females per generation in both breeds. However, such an increase was much faster in the Gasconne, which, nonetheless, reached a total number of breeding individuals/generation lower than that of the Barbezieux (Figure S10).

### Effective population size

As expected from the larger size of the founding nucleus, *N*_*e*_ was estimated at 88.48 (90% CI: 66.97 - 116.89) in the Barbezieux, as compared to that of the Gasconne, which was estimated at 69.07 (90% CI: 53.24 - 89.62).

### Temporal changes in deleterious variation

We have shown that since the start of the conservation program genetic diversity declined (Δ*π*) at the costs of an increase in realised (Δ*F*_*ROH*_) and expected (Δ*F*_*ped*_) inbreeding, resulting from an accumulation of longer ROH (≥ 3 Mb), particularly in the Gasconne breed. To verify whether the decline in Δ*π* and the increase in Δ*F*_*ROH*_ and Δ*F*_*ped*_ were associated with changes in deleterious variation, we annotated variants with respect to their predicted impact on the encoded amino-acid into benign (synonymous and tolerated missense mutations) and damaging (deleterious missense mutations and mutations that disrupt splice sites or start or stop codons).

We first looked at the derived allele frequency (DAF) spectrum to examine the impact of purifying selection on our samples. The two breeds had more fixed, high frequency (DAF >0.90) benign (i.e., synonymous, tolerated) mutations than (putative) damaging (i.e deleterious, LoF) ones at the same frequency (Figure S11). To determine how common purging was in the two breeds, we looked at two measures of genetic load (L) (Fig. 4). In both breeds we observed a net increase in GERP load (Fig. 4a) and a reduction in chCADD load (Fig. 4b) in the period 2003 - 2013/15. The increase in GERP load was significant only in the Gasconne (Wilcox test *p-value*: 0.0094), as well as the reduction in chCADD load (Wilcox test *p-value*: 0.0327).

**Figure 4.**
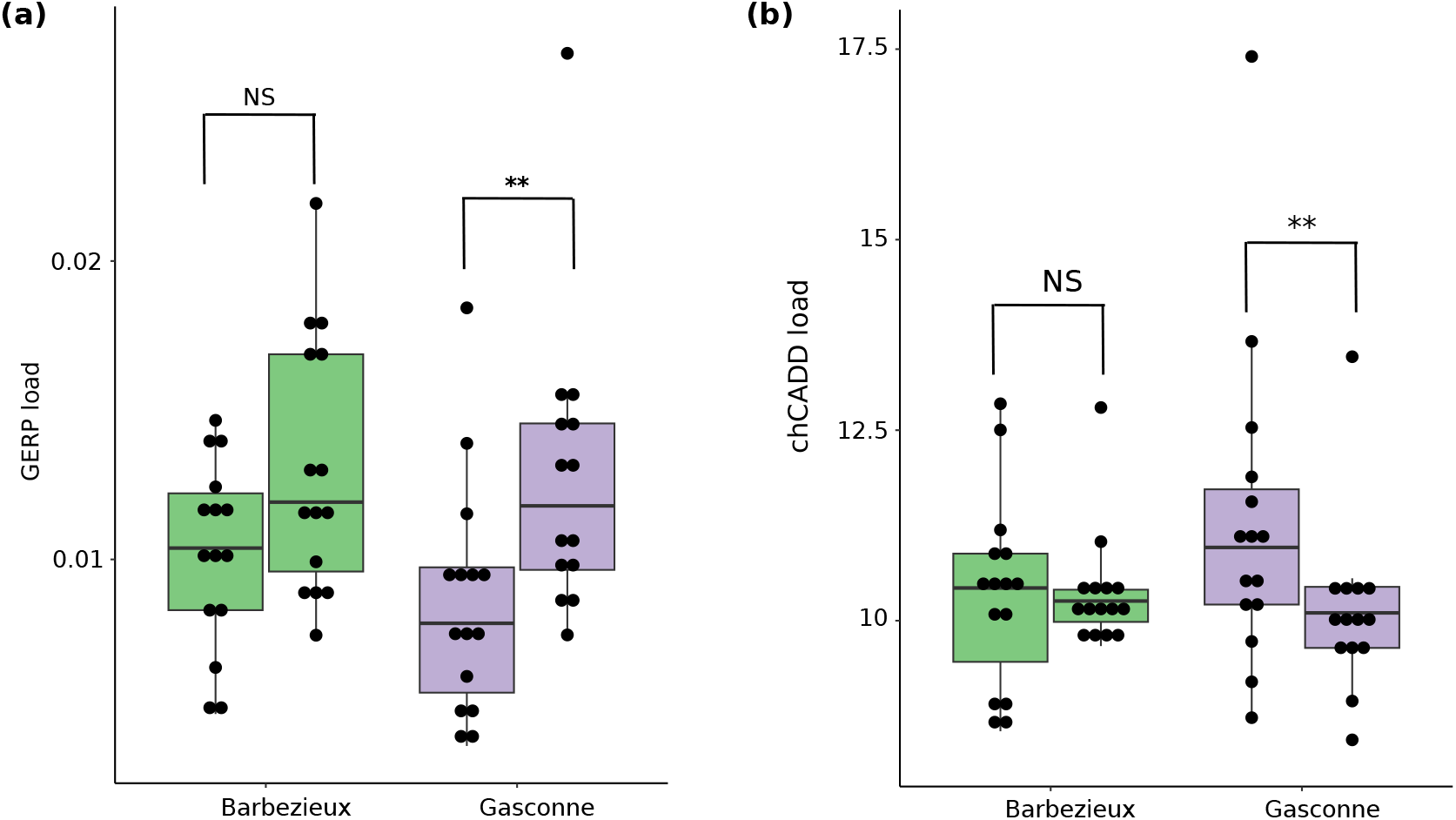
Genetic load in the Barbezieux and Gasconne breed. **a**. Genetic load approximated using the GERP score information of each homozygous damaging mutation (GERP > 1.0). **b**. Genetic load approximated from all variants independently of their functional annotation using the chCADD score.

### Signatures of positive selection

The conservation program here studied was established with the objective of exploiting local breed’s diversity for the production of products under quality labels. Hence, positive selection was expected to some extent. To test this hypothesis, we identified genomic regions under selection (or selective sweeps) using the HMM approach developed by Paris et al. (2019). We decided to perform this analysis only on the Barbezieux, as the pedigree data of the Gasconne did not make it possible to relate the two sets of animals sampled. After filtering SNPs for an FDR threshold of 5%, no significant SNPs were identified, meaning that positive selection, if it occurred over the 10 generations, was weak enough to not leave any detectable signature in the genome.

This result was further supported by the allele frequency distribution, which remained unchanged in the 10 generations (Figure S12).

### Phenotypic data: productive and reproductive performance

We analysed one productive (Table S5, Table S6) and 6 reproductive traits (Table S7, Table S8) collected and provided by the SYSAAF to look at possible changes in productive and reproductive performance over the 10 generations. In the Barbezieux, it seems that most of the selection effort for the trait body weight at 8 weeks took place between 2004 and 2006, where body weight was higher than in the founder generation, reaching almost a value of 1,200g (Figure S13). However, after 2007 body weight decreased to below 1,000g, to then increase once again between 2008 and 2010 and between 2011 and 2013. While over the 2003-2013 period the trait body weight at 8 weeks had in the Barbezieux an overall positive trend with some initial fluctuations, we did not observe any clear phenotypic trend in the Gasconne breed over the 2010-2015 period (Figure S13). Regarding reproduction, the % fertile eggs and the % hatched eggs increased in the Barbezieux breed, leading to a positive selection coefficient in the individuals sampled in 2013 (Table S7). Late embryonic mortality remained rather constant in the same time period (Table S7). Thus, the total number of chicks hatched increased because fertility had increased.

The situation in the Gasconne was quite difficult to analyze since reproductive data were only available for the 2013 generation (Table S8), making any prior trend estimate impossible. However, when looking ahead (2013-2018), we found that all six reproductive traits have a fluctuating trend, suggesting difficulties in management and the absence of clear selection objectives.

## Discussion

In this era of rapid decline in biological diversity, conservation programs have become critical for preserving the genetic diversity harbored by individual genomes (Kleinman-Ruiz et al., 2019). The importance of a conservation program on a species genome has extensively been addressed in endangered wild species (Kleinman-Ruiz et al., 2019; Robinson et al., 2019; Van Der Valk et al., 2019; Xue et al., 2015), but in local livestock breeds this has rarely been done. The rationale is that, when in place, management of local breeds cannot afford the cost of collecting high-density SNP data or, even less likely, whole-genome sequencing data. This study represents a unique case in Europe of local chicken breeds under a conservation program. Whereas the studied breeds already had SNP genotyping data for a single generation, this study is one of the few where temporal whole-genome sequencing data were used as a tool to gather critical information on the demographic and genetic processes accompanying a conservation program, with the ultimate goal of informing management and aid decision-making to keep local breeds from the brink of extinction.

We are aware that one of the limitations of this study is the sample size, as only 15 samples were selected and analysed for each time period (2003 and 2013). However, this range of magnitude is similar to the number of founder animals used for each breed and can be considered as representative of the initial sampling. Furthermore, such sample size is not uncommon in studies of local livestock breeds and endangered species, for which the availability of hundreds of individuals is unachievable due to their small population size. Finally, by sequencing the entire genome of each of the 15 samples included in the study, we believe we were nonetheless able to estimate key demographic parameters more reliably than by means of high-density SNP chip arrays on a larger number of individuals.

### How to assess the success of a conservation program

In this study, we showed that the conservation and exploitation of local breeds diversity is a valuable strategy as it allows dynamic breed conservation. The conservation program of the Barbezieux and Gasconne was similar in organisation and set up to that of the Bresse, a local chicken breed native to the homonymous province in eastern France (Verrier et al., 2005). Similar to the Bresse, members of the founding nucleus were sampled from fancy breeders in the geographical area of origin of the breed, which is often defined by law. Moreover, only individuals complying with the phenotypic standard were qualified by the SYSAAF to establish the selection line at the Centre de Sélection de Béchanne. The conservation program of the Bresse has shown that when a product becomes a success, the risk status of a breed can be improved, while the loss of a breed’s specific abilities can be prevented. Although in its infancy, the conservation program of the Barbezieux and Gasconne aims to achieve a similar success by linking the name of a breed to a product that has a controlled designation of origin status. Despite this, the analysis of the productive and reproductive traits suggests that in both breeds more emphasis was placed, for now, on the maintenance of the breed’s standards rather than on the selection for enhanced productive traits (for example, body weight). Nonetheless, we found the Barbezieux breed to be slightly more productive than the Gasconne, despite the loss of heterozygosity and increase in inbreeding over the 10 years. It is possible that in the Barbezieux the slightly higher productivity is the result of genetic purging taking place following the increase in inbreeding, which reduced the number of functionally deleterious mutations genome-wide (Fig. 4b). However, it is also possible that this higher productivity is the result of historical selection for more productive animals.

The lack of detection of selective sweeps may also be due to the mild selection intensity that was applied. Selection pressure was all the more limited that the number of sires and dams has been increasing for the Barbezieux breed which suggests that adult fertility was a key parameter. Indeed, fertility has improved, but it could be for management reasons as well as for genetic reasons. Although our selective sweep analysis failed at identifying genomic regions under positive selection, we cannot rule out the possibility that a mild form of selection has nonetheless been taking place.

### The importance of management in a conservation program

From a genetic standpoint, the primary objective of a conservation program is to maintain the highest possible levels of genetic diversity, while controlling for the increase in inbreeding. By doing so, populations will be able to respond to future changes in breeding goals and avoid a reduction in fitness (De Cara et al., 2013). As our analyses on the pedigree data clearly illustrate, conservation programs are generally founded by a small number of individuals, often coming from breeds that have a small population size themselves. Therefore, the first step in safeguarding genetic diversity is to capture as much variation as possible in the founding nucleus. This was not really the case here, where a very small number of founders were chosen for each breed. The second step is the genetic screening of the founding nucleus. Individuals selected for a conservation program may carry several genetic risks including (1) low genetic diversity, (2) high level of inbreeding, and (3) accumulated deleterious alleles. The most common practice to mitigate these genetic risks in a conservation program is the minimisation of average kinship (Caballero and MA Toro, 2000; BJ Fernández and M Toro, 1999; Meuwissen, 1997), which was applied to the Barbezieux since 2003 and to the Gasconne since 2009. According to this strategy, the control over inbreeding (or co-ancestry) can be achieved if each individual contributes to the next generation with an optimal number of offspring (De Cara et al., 2013; Meuwissen, 1997). Hence, the effective population size (*N*_*e*_) is maximised (Meuwissen, 1997), while the expression of (recessive) deleterious mutations is minimised. However, in order to implement this management strategy, information on individual relationships is required (De Cara et al., 2013), which is not trivial both in domesticated and wild species, but was available in the present study. The minimum kinship strategy implemented by the SYSAAF is based on the traditional analysis of pedigree data for the selection of breeding individuals. The effects of the average kinship strategy were particularly visible in the *F*_*ped*_ values of the Barbezieux, which, after an initial steep increase, stabilized at around 7%. Molecular data enabled us to take a step forward in the analysis of inbreeding, allowing us to separate the past from recent inbreeding. As a result, we were able to show that mating between close relatives (recent inbreeding), which is exemplified by the accumulation of longer autozygous segments, should be avoided as much as possible in future breeding decisions. In the case of the Gasconne, although recent inbreeding is of major concern, we cannot exclude that the recent bottleneck associated with the establishment of the conservation program in 2009 may have contributed as well. Our findings illustrate two important aspects. First, that if management is properly carried out (i.e., Barbezieux), a conservation program can still thrive even when established from a small number of founders. Hence, resampling of individuals should be carefully evaluated to limit any negative effects of changes in management on animal genetic diversity. And second, that whenever possible, pedigree information should be recorded to elucidate management.

### The role of a conservation program in purging deleterious mutations

As our study shows, in conservation programs where the population is treated as a closed nucleus, inbreeding can rapidly increase along with the probability of exposing deleterious alleles in homozygous state. A common mitigating strategy designed to restore genetic diversity and reduce inbreeding is the introduction of new individuals (and genes) from a source into a target population. In the case of the Barbezieux and Gasconne, introduction of genetic material from other breeds (introgression) is highly discouraged to preserve the genetic uniqueness of the breed. Therefore, introduction of genetic material from individuals of the same breed could offer a valuable solution to the observed loss of genetic diversity and increase in (genomic) inbreeding. High-throughput sequencing data can guide this decision, as they provide additional information on the often-neglected functional relevance of variants (Bosse, Megens, Madsen, Crooijmans, et al., 2015; Oosterhout et al., 2022).

Deleterious mutations have important consequences on an individual’s survival and genetic potential. Conservation programs established without the support of molecular data are very likely to retain deleterious mutations, reducing, in the long-term, population mean fitness (De Cara et al., 2013). Deleterious mutations are a valuable source of information to perform *in-silico* prediction of fitness. Compared to previous studies that focused on protein-coding variants (Bortoluzzi, Bosse, et al., 2020; Bosse, Megens, Derks, et al., 2019; Derks et al., 2017), we here estimated genomic fitness genome-wide by focusing on all mutations independently of their coding potential. Such major breakthrough is now possible thanks to the development of the ch(icken) CADD model (Groß et al., 2020), an integrative annotation tool that can effectively score and prioritise variants genome-wide. Our findings on the genomic fitness suggest that, in the case of the Barbezieux, introduction of genetic material from individuals outside the nucleus would be beneficial for the long-term conservation of the breed. However, for this management practice to succeed, individuals chosen to genetically rescue the current population should be functionally screened along with the members of the nucleus by either whole-genome sequencing data or a high-density SNP chip specifically designed for this purpose. This screening procedure should not be underestimated, as large populations with high genetic diversity may harbor recessive deleterious alleles that, if introduced in a small population, could put this population at higher risk of extinction (Bertorelle et al., 2022). While introduction of genetic material might help restore the genetic diversity in the Barbezieux, the impact of this strategy on the Gasconne is difficult to predict due to the different genetic make-up of the 2003 and 2013 founding population. We therefore recommend the SYSAAF to sequence individuals belonging to the 2009 founding nucleus in the coming years to better monitor changes in genetic diversity, inbreeding, and genomic fitness.

### The added value of whole-genome sequence data to assess the conservation status of a population

The Barbezieux and Gasconne breeds were included in a large-scale study aimed at comparing various indicators of genetic diversity of local chicken breeds on the basis of 57K SNP genotyping of one generation in 2013 (Restoux et al., 2022). Both breeds exhibited very similar values for all indicators that are commonly calculated (*F*_*it*_, *F*_*is*_, *H*_*o*_, *H*_*e*_, MAF, fixed alleles) and slightly different values for *F*_*ROH*_ with a higher value for the Gasconne breed, as confirmed here. Here we show that whole-genome sequence data were much more efficient than SNP genotyping to reveal the differences between the two breeds. For instance, whole-genome sequence data allowed us to estimate the effective population size (*N*_*e*_) of both breeds and changes in selection over time. At the same time, the higher number of markers sampled across the genome allowed us to estimate changes in genetic load over the 10 years time, which we here estimated from both protein-coding and non-coding SNPs. Another advantage of using whole-genome sequence data over SNP chip data is due to the ascertainment bias SNP chip data are prone to, in particular in local breeds, which are often not used to design the SNP chip panel. Ascertainment bias can result in biased estimation of diversity metrics, while whole-genome sequence data are less prone to such artifacts, especially when variants are polarised. The higher resolutive power of sequencing data is expected, but the present results show that generation of whole-genome sequence data should be planned at regular intervals to better monitor the genetic status of a conserved breed while looking at parameters (e.g., genetic load) that often go unchecked but that can impact the performance of future generations. Sequencing costs are decreasing and accumulating such data could later on make it possible to impute whole-genome sequence from SNP array data also for local livestock breeds.

### *Ex situ* conservation practices in domestic animal diversity

In the context of domestic animal diversity, *ex situ* conservation practices are recognised as an essential complementary activity to *in situ* conservation actions for the maintenance of a broader genetic base. In this study, the conservation program of the Barbezieux and Gasconne relies on the maintenance of live animals (i.e., *in vivo*), though cryoconservation (i.e., *in vitro*) has been performed for one generation sampled along the program. As gene bank collections are stored for an indefinite time, they allow to preserve genetic diversity from demographic and genetic forces, such as selection and genetic drift. The interest for cryopreservation has increased over the years also for local livestock breeds, and specifically for poultry, thanks to the development of reproductive biotechnologies and efforts to enhance the use and exploitation of genetic collections (Blesbois et al., 2007). Although a gene bank is in most cases regarded as a safety collection and a complement to *in situ* and *ex situ in vivo* conservation programs, stakeholders directly involved in conservation efforts should also take advantage of existing national gene banks to regularly store genetic material for use in the future. This is particularly relevant for local breeds as their small size puts conservation programs at higher risks of failure if not properly managed and supported by molecular data, as this study shows. In the case of the Barbezieux, we encourage the analysis of the genetic material stored in the gene bank, since it may be used to reintroduce lost diversity. In the case of the Gasconne, the semen stored after 2009 is likely insufficient to reintroduce diversity. Hence, the sustainability of the conservation program would benefit, once again, from additional sequencing.

## Acknowledgements

The authors would like to thank Cyriel Paris and Simon Boitard for their support on the use of the Hidden Markov Model for the selection signature analysis. We would also like to thank the technicians and scientists from the Centre de Sélection de Béchanne and SYSAAF for the collection and sharing of data. We are grateful to Gilbert Marchand and Nicole Billion for the Barbezieux breed and to Thierry Dubarry, Elodie Menvielle and Sylvie Blagny for the Gasconne breed for their collaboration in providing data and explanations regarding the history of the breeding programs. A special thanks to the breeders for their cooperation during data collection. And last, but not least, a special thanks to the two reviewers, Markus Neuditschko and Claudia Fontsere Alemany, for their valuable comments, which we believe have enhanced the quality and relevance of our work.

## Author contributions

M.T.B conceived the study and organised the data collection with the breeders and SYSAAF. C.B. designed the study, carried out the genomic analyses, and wrote the manuscript. R.R., B.D., and F.P. collected and provided the pedigree and phenotypic data. G.R., M.B., and M.T.B jointly supervised the study and contributed to the writing of the manuscript. All authors revised and approved the final version.

## Data, script, code, and supplementary information availability

All the scripts necessary to run the analyses presented here can be found in Zenodo: https://doi.org/10.5281/zenodo.12744831. All ch(icken) CADD (chCADD) scores are publicly available in the Open Science Framework (OSF) at https://osf.io/8gdk9/. The raw data of the 59 individuals sequenced for this study are archived in the European Nucleotide Archive (ENA) under project number PRJEB72503. All data needed to evaluate the conclusions in the paper are present in the paper and/or the Supplementary Materials, which can be found in Zenodo: https://doi.org/10.5281/zenodo.12912874.

